# A Yeast-Based High-Throughput Screening Platform for the Discovery of Novel pre-mRNA Splicing Modulators

**DOI:** 10.1101/2025.10.20.683303

**Authors:** Sierra L. Love, Henrik Vollmer, Ya-Chu Chang, Joshua C. Paulson, Tucker J. Carrocci, Melissa S. Jurica, Hai Dang Nguyen, Aaron A. Hoskins

**Author notes:** CORRESPONDING AUTHOR: Aaron A. Hoskins.

## Abstract

Pre-mRNA splicing is a crucial process in eukaryotic gene expression, and splicing dysregulation has been linked to various diseases. However, very few small molecules have been discovered that can modulate spliced mRNA formation or inhibit the splicing machinery itself. This study presents a novel high-throughput screening (HTS) platform for identifying compounds that modulate splicing. Our platform comprises a two-tiered screening approach: a primary screen measuring growth inhibition in sensitized in *S. cerevisiae* (yeast) strains and a secondary screen that relies on production of a fluorescent protein as a readout for splicing inhibition. Using this approach, we identified 4 small molecules that cause accumulation of unspliced pre-mRNA *in vivo* in yeast. In addition, cancer cells expressing a myelodysplastic syndrome-associated splicing factor mutation (SRSF2^P95H^) are more sensitive to one of these compounds than those expressing the wild-type version of the protein. Transcriptome analyses showed that this compound causes widespread changes in gene expression in sensitive SRSF2^P95H^-expressing cells. Our results demonstrate the utility of using a yeast-based HTS to identify compounds capable of changing pre-mRNA splicing outcomes.

## INTRODUCTION

Pre-mRNA splicing, the process of removing introns and joining exons to produce mature mRNA molecules, is a critical regulatory step in eukaryotic gene expression. This essential process is orchestrated by the spliceosome, a large ribonucleoprotein complex composed of five small nuclear RNAs (snRNAs)—U1, U2, U4, U5, and U6—each associated with proteins to form small nuclear ribonucleoproteins (snRNPs). Precise splicing is crucial for ensuring the production of functional proteins that drive cellular function and organismal development. Dysregulation of pre-mRNA splicing, whether through mutations in spliceosome components or alterations in RNA substrates, has been linked to a broad spectrum of human diseases including cancers, neurological disorders, and rare genetic syndromes^1^.

Despite pre-mRNA splicing being studied for decades, there are few ways to chemically modulate or inhibit splicing *in vivo* by targeting the splicing machinery itself^2^. This is especially apparent when compared with the chemical tools available to inhibit protein translation at numerous distinct stages^3^. Relatively few analogous chemical probes of splicing mechanism exist. While it is possible to fuse tags to protein splicing factors to change their activity or abundance in response to a small molecule (*e.g.,* degron tags)^4^, only a handful of molecules have been discovered that can change splicing by direct interaction with the endogenous machinery. The most widely used splicing inhibitors are small molecules that target the U2 snRNP protein SF3B1. These compounds (including pladienolide B, herboxidiene, and spliceostatin derivatives) are potent inhibitors of splicing in humans by binding (sometimes covalently) to a protein pocket on SF3B1, near its interaction site with the PHF5A protein^5^. The PHF5A protein can form a covalent adduct with some of these inhibitors, leading to irreversible inhibition^6^. This pocket normally accommodates the U2 snRNA/branch site intron RNA duplex, which cannot bind correctly if the drug is present. Even though SF3B1 is overall highly conserved, yeast SF3B1 (Hsh155) is naturally resistant to inhibition. Substitution of human for yeast Hsh155 protein domains or mutation of the drug binding site in Hsh155 confers drug sensitivity^7, 8^. These mutated yeast strains have subsequently been used for studies of splicing and intron structure due to their susceptibility to specific and rapid splicing inhibition^9, 10^. A few other splicing inhibitors have been discovered but are much less widely used and characterized and/or have controversial mechanisms of action^11^.

In addition to their utility in laboratory settings, small molecule modulators of splicing can show considerable clinical promise. This also has best been studied in terms of SF3B1-binding compounds that have been used to selectively kill cancer cells with mutations in splicing factors, such as those found in hematological malignancies^12, 13^. It is thought that dysregulated splicing, accumulation of intronic RNA in the cytosol^13, 14^, and other RNA metabolism abnormalities render these cells uniquely sensitive to splicing modulation. This can create a therapeutic window that spares normal cells^15, 16^. Alternatively, these drugs can be used to induce formation of novel protein isoforms (proteoforms) via alternative splicing of transcripts, and the drug-dependent proteoforms can in turn be used as cancer cell markers or for targeted neoantigen-based therapy^17^.

In this study, we developed and validated a *Saccharomyces cerevisiae* (yeast)-based high-throughput screening (HTS) platform for identifying small molecules that can change splicing outcomes. We identified four compounds that function as splicing modulators *in vivo* in *S. cerevisiae*. K562 cancer cells expressing the SRSF2^P95H^ splicing factor mutation are selectively sensitive to one of these compounds, and drug treatment results in changes in gene expression and splicing outcomes. Together, these findings highlight the utility of a yeast-based HTS for finding new small molecule modulators of pre-mRNA splicing for potential biochemical and medical applications.

## MATERIALS AND METHODS

### Chemicals and Drug Libraries

Drug libraries include: Selleck Chem FDA-approved drugs, RIKEN-Pilot Natural Products Depository (NPDepo), Life Chemicals 4 (LC4), and National Cancer Institute’s (NCI) Developmental Therapeutics Program (DTP) and Experimental Program (NExT) Diversity libraries. Phleomycin and herboxidiene were diluted in water and DMSO, respectively, then aliquoted and stored at -20°C. The chemical libraries were diluted in DMSO to a final concentration of 10 mM and stored at -80°C.

### Preparation of Plasmids and Yeast Strains

Wild-type (WT) and humanized Hsh155 plasmids and corresponding yeast strains have been previously described^7, 8^. The plasmid BRR2 pRS313 (Addgene 111411; Guthrie lab) was modified to remove three EcoRI sites, one SacI site, and add an AflII site using Quikchange Lightning Multi (Agilent). These modifications facilitated further plasmid alterations without causing amino acid changes in Brr2. The resulting plasmid (pAAH1347) is referred to as wild type (WT). Plasmid pAAH1459 was created by PCR amplification of pAAH1347 to produce a 10.8 kb linear fragment using Herculase II (Agilent). This fragment was combined with a synthetic gBlock encoding humanized Brr2 regions using NEBuilder HiFi. The exchanged regions spanned from amino acids L1202 to L1300 and L1540 to V1741. To construct Brr2 strains for plasmid shuffling and screening, three ABC transporters (PDR5, YOR1, SNQ2) were sequentially deleted from strain BY4741 using a CRISPR-based approach^18^, resulting in the removal of the entire ORF of each transporter without cloning scars or genomic incorporation of resistance markers. The resulting strain (yAAH3007) was transformed with either pAAH1347 or pAAH1459, and transformants were selected on synthetic dropout (SD) media lacking uracil (SD-URA). The genomic copy of BRR2 was replaced with the hphMX6 gene cassette encoding hygromycin resistance through homologous recombination^19^.

Reporter plasmids containing genes for fluorescent proteins with introns were generously provided by the Manny Ares lab (UCSC) and based on the modular yeast toolkit for gene assembly^20^. Briefly, the plasmid constructs contain a kanamycin resistance cassette, URA3 marker, and URA3 homology regions flanking a reporter gene driven by the PGK1 promoter. This produces an RNA transcript with the PGK1 5’ UTR, an open reading frame (ORF) encoding an N-terminally FLAG and 6xHis-tagged yellow fluorescent protein (Venus), and the ADH1 3’ UTR. The ORFs were interrupted with introns based on the first intron of MATa1. Two different reporters were used: (1) the splicing-in-frame (SPLIF) reporter in which the splicing of the pre-mRNA removes the intron to leave a mRNA that properly codes for the Venus protein and (2) the splicing-out-of-frame (SPLOOF) reporter in which intron removal causes a change in the reading from the protein and lack of Venus production.

Plasmids were linearized with NotI to allow integration at the URA3 locus and selection on -URA dropout plates. Linearized DNA was purified by agarose gel electrophoresis [1% (w/v) low-melt agarose] followed by a Promega SV Gel and PCR Cleanup kit to remove the DNA from the agarose. The linear DNA fragment was then transformed into yeast strains using standard techniques^21^ and selection carried out by plating on -URA DO plates. Correct integration was confirmed by PCR and sequencing of the gene isolated from the modified yeast strains.

### Primary Screening of Drug Libraries

Screening was conducted at the University of Wisconsin Carbone Cancer Center Small Molecule Screening Facility (SMSF). Controls (DMSO, herboxidiene, and phleomycin) and library compounds were dispensed into 384-well clear plates (Greiner Bio-One) using an Echo550 liquid handler.

Yeast strains were cultured overnight in DO-Trp (for Hsh155 strains) or DO-His (for Brr2 strains) liquid media with shaking at 220 rpm at 30°C. Before screening, the cells were diluted to an optical density at 600nm (OD_600_) of 0.1 and then grown to log phase (OD_600_ ∼1.0) under the same conditions. The culture was then diluted to an OD_600_ of 0.0075 in low nitrogen media made by using yeast nitrogen base lacking both amino acids and ammonium sulfate and by addition of 1 g/L monosodium glutamate along with the appropriate mix of amino acids and nucleotides to select for growth^22^.

The diluted yeast were then added to 384-well plates (50µL per well; Greiner Bio-One) using a BioTek Microflo Select liquid handler. Cells were incubated in a shaking incubator (220 rpm at 30°C) for 24 h. After incubation, cells were centrifuged at 1000g for 1 minute to remove air bubbles and shaken at 2000 rpm for 4 min to ensure uniform readings in each well. Plates were then read using a PHERAstar (BMG LABTECH) plate reader.

The Bland-Altman approach^23^ was used to assess the reproducibility of yeast growth between replicates. In summary, the differences between replicates and the mean growth values were calculated for each well. These values were then plotted, and the upper and lower 95% confidence limits were determined to represent the maximum variability between replicates. Analysis of normalized growth data indicated that the differences between yeast growth replicates ranged from -0.2 to 0.1, demonstrating that growth was highly consistent across replicates.

The screen quality was evaluated by calculating the statistical Z’ values, with Z’ values between 0.5 and 1.0 indicating a robust, high-quality assay ^24^. All screenings resulted in plate Z’ values between 0.70 and 0.95. Collected data were uploaded to the Collaborative Drug Discovery (CDD) database for quality control and normalization. Each drug library, except for the RIKEN NPDepo and the NCI DTP, was screened once per strain, and compounds showing at least 30% growth inhibition relative to the DMSO controls were then re-screened in duplicate (for a total of three replicates for these compounds). The RIKEN library was screened in triplicate for all strains, and the NCI DTP was screened in duplicate for all strains.

### Secondary Screening of Compounds that Inhibit Yeast Growth

Yeast strains with integrated SPLIF or SPLOOF reporters were grown overnight in DO-Trp or DO-His liquid media at 30°C with shaking (220 rpm). Cells were then diluted to an OD_600_ of 0.1 in their respective dropout media. A total of 99 µL of these cultures was combined with 1 µL of DMSO or a library compound dissolved in DMSO in a Corning Costar 96 or 384-well black clear-bottom cell culture plate. All compounds were at a final concentration of 10 µM. The plates were covered with Breathe-Easy® plate sealing membranes to minimize evaporation and then incubated at 30°C with shaking at 220 rpm for 24 hours in a Tecan Infinite M1000Pro plate reader. Turbidity (OD_600_) and fluorescence intensity (512 nm excitation, 535 nm emission) were measured every 15 min. For data analysis, each condition was normalized to the DMSO control. Lead candidates were identified as those that reproducibly increased normalized fluorescence intensity (> 1.0) for the SPLOOF reporter and decreased intensity for the SPLIF reporter (< 1.0) for at least ten consecutive cycles (∼2.5 h). These screens were performed in duplicate for each compound.

### RT-PCR to Validate pre-mRNA Accumulation

Yeast cell cultures were grown overnight in SD-Trp or SD-His liquid media and then diluted to an OD_600_ of 0.5. Cells were then treated with each drug at the indicated concentration (see below) while shaking at 220 rpm at 30°C. After 10 h, samples were collected and pelleted. RNA was extracted from yeast cells using the NucleoSpin® RNA extraction kit (Machery-Nagel). Splicing of the endogenous MATa1 transcript was then assessed by reverse transcription and PCR using RNA (150 ng) isolated from each sample and Promega Access RT-PCR System kit in 25 µL reactions. RT-PCR results were analyzed using agarose (2% w/v) gel electrophoresis to separate spliced and unspliced bands using 3-4 biological replicates. Gels were photographed and images analyzed using the ImageQuant analysis toolbox (Cytiva). Band intensities were quantified to calculate the percent spliced for each reaction (mRNA/(pre-mRNA + mRNA)). Statistical analyses (student’s *t*-test) were performed using Excel and GraphPad Prism.

### *In Vitro* Splicing Reactions

Assays were conducted as previously described ^25^. In brief, nuclear extracts (NE) were prepared from HeLa cells grown in DMEM/F-12 1:1 and 5% (v/v) newborn calf serum. [^32^P]-radiolabeled G(5′)ppp(5′)G-capped AdML pre-mRNA substrate was generated by T7 run-off transcription followed by denaturing polyacrylamide gel purification. Each reaction consisted of potassium glutamate (60 mM), magnesium acetate (2 mM), ATP (2 mM), creatine phosphate (5 mM), tRNA (0.1 mg/ ml-1), HeLa nuclear extract (40% v/v), and pre-mRNA substrate (2 nM). Reaction reagents were incubated with DMSO (1% v/v) and each drug at the corresponding concentrations for 60 min at 30°C. RNA was extracted using phenol-chloroform, precipitated using ethanol, dissolved in deionized formamide, and then resolved on a denaturing acrylamide gel (7 M Urea, 15% (w/v) acrylamide).

### Cell culture

The *SRSF2^P95H^* mutation was introduced into its endogenous locus in K562 cells using CRISPR-Cas9, generating *SRSF2^P95H/+/+^*cells. SRSF2^WT^ and SRSF2^P95H^ mutant K562 cells were maintained in IMDM medium (Gibco, 12-440-079) supplemented with penicillin–streptomycin (100 U/mL, Gibco, No. 15140122), GlutaMax (1%, Gibco, No. 35050061), and cultured in a 37°C/5% CO_2_ incubator.

### Cell Viability Assays

K562 SRSF2^WT^ and SRSF2^P95H^ isogenic cells were seeded in white flat-well 96-well plates at a density of 200-300 cells per well^26^. Cells were treated with compounds (0-50 µM) or a DMSO control for 7 days. Cell viability was subsequently determined using CellTiterGlo (Promega, Cat. G7571) according to the manufacturer’s instructions. The proportion of viable cells with drug treatment was calculated relative to the DMSO control. A four-parameter non-linear fit of inhibitor concentration vs. response was performed in GraphPad Prism v10.0 (GraphPad Software, San Diego, CA; RRID:SCR_002798).

### RNA-seq Sample Preparation and Sequencing

K562 SRSF2^WT^ and SRSF2^P95H^ isogenic cells (500,000 per condition) were seeded and treated with 16.7 µM of C7 for 24 h. Total RNA was isolated using TRIzol® according to the manufacturer’s instructions (Zymo Research, Cat. R2072) with DNase I treatment. Quantification was performed using a Qubit Flex Fluorometer (Thermo Fisher Scientific). RNA quality control, library preparation, and sequencing were performed by GENEWIZ (Azenta Life Sciences). Library prep involved poly-(A) selection, cDNA synthesis, and adapter ligation. Sequencing was completed on an Illumina® NovaSeq^TM^ platform with a target depth of ∼50-70 million paired-end 150 bp reads per sample. Data are available via the NCBI Sequence Read Archive (BioProject ID PRJNA1271094).

### RNA-seq Data Analysis

Sequence reads were trimmed to remove adapter sequences and nucleotides with poor quality using Trimmomatic v0.36. The trimmed reads were mapped to the *Homo sapiens* GRCh38 reference genome available on ENSEMBL using the STAR aligner v2.5.2b. Unique gene hit counts were obtained using featureCounts from the Subread package v1.5.2. GENEWIZ (Azenta Life Sciences) performed trimming, alignment, and gene count quantification. Differential gene expression analysis was conducted with DESeq2 v1.46.0 ^27^. Genes were considered differentially expressed (DEG) if they met the thresholds of log_2_ fold change > 1 or < -1 and an adjusted *p*-value < 0.05. Alternative splicing (AS) analysis was performed using rMATS^28^. Differentially spliced events were identified using a percent spliced-in (ΔPSI) change > 0.1 (10%) and a false discovery rate (FDR) < 0.05. Functional enrichment analysis of significant DEGs and alternatively spliced genes was conducted using ShinyGo with enriched Gene Ontology (GO) terms identified against the human genome background^29^.

## RESULTS AND DISCUSSION

### Primary Screen Development and Optimization

To identify molecules that act as splicing modulators, we first aimed to select compounds that result in growth inhibition of yeast since pre-mRNA splicing is essential for viability. We developed a primary screen to assay growth in 384-well plates in which the growth endpoint was measured after 24 h (**Fig. 1B**). In order to bias for selection of splicing modulators, we utilized yeast strains containing variant, non-native splicing factors. We chose to screen strains containing either WT or “humanized” genes for Hsh155/SF3B1 or Brr2 proteins (**Fig. 1C**).

**Figure 1.**
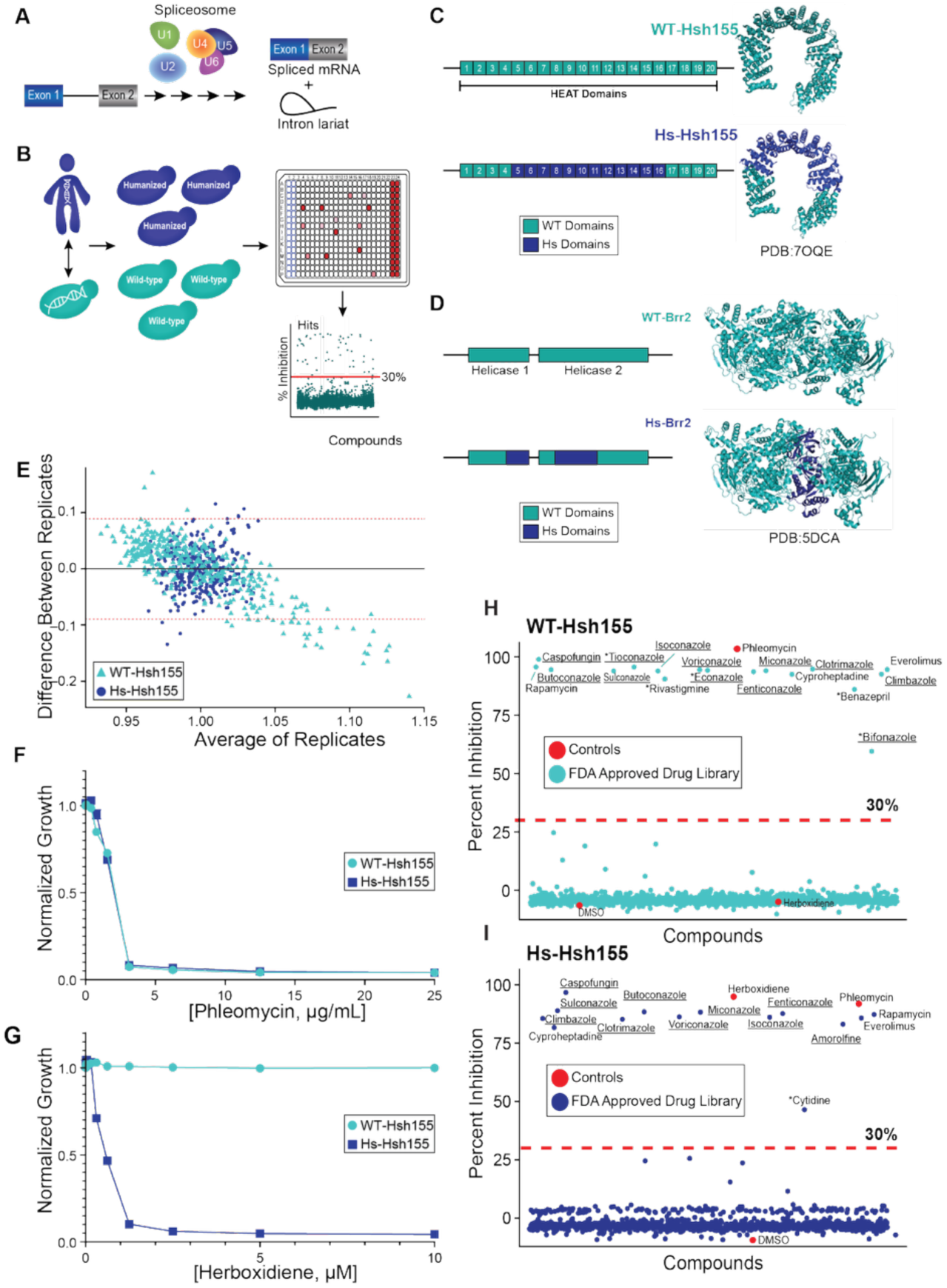
Primary Screen Development and Optimization. **(A)** Overview of pre-mRNA splicing: introns are removed and exons joined to generate mature mRNA. This process is catalyzed by the spliceosome, composed of five small nuclear ribonucleoproteins (U snRNPs). **(B)** Schematic of primary screening strategy. Compounds were screened in 384-well plates using wild-type and humanized yeast strains. Hits were identified as those that reduced yeast growth by ≥ 30% relative to DMSO controls. **(C, D)** Diagrams of chimeric Hsh155 and Brr2 proteins used in engineered yeast strains. **(E)** Bland-Altman plot assessing reproducibility between biological replicates. Red dotted lines indicate 95% confidence limits. Both the x- and y-axes are expressed in terms of absorbance (OD_600_) units. **(F, G)** Dose-response curves of controls (phleomycin and herboxidiene) in WT and Hs-Hsh155 yeast. Relative growth was measured at 24 h post-treatment and normalized to DMSO controls. **(H, I)** Pilot screen of Selleck Chem FDA-approved drug library. Percent growth inhibition of Hsh155 strains relative to DMSO controls are shown; red dotted lines indicate 30% inhibition thresholds. Antifungals are underlined; strain-specific hits are marked with asterisks.

We focused on these proteins since they have already proven susceptible to drug targeting in human cells^15, 30, 31^. We reasoned that use of these strains might lead to identification of compounds that modulate splicing due to sensitization from lack of the endogenous splicing machinery. Alternatively, this could lead to potential identification of compounds that target SF3B1 or Brr2 proteins directly. Further, we have already shown that yeast-human chimeras of Hsh155/SF3B1 confer sensitivity to splicing modulators and growth inhibition, allowing us to use these strains and compounds as positive controls during development of the screening methodology. In the case of Hsh155/SF3B1, the protein was humanized by exchange of heat repeats 5-16 of the yeast protein for the corresponding human domains (**Fig. 1C**) as previously described^7, 8^. Brr2’s domain organization consists of two helicase cassette modules. In this case, we humanized the domain interface between these modules (**Fig. 1C**). This region is less well-conserved between yeast and humans and is the binding site for allosteric inhibitors of Brr2 activity^30^. Finally, we utilized yeast strains lacking multi-drug efflux pumps and grew yeast in low-nitrogen media in order to maximize sensitivity to compounds^8, 22, 32^.

We next optimized yeast growth conditions in the high-density 384-well plates. In preliminary studies, we grew humanized (Hs-Hsh155) and WT Hsh155 (WT-Hsh155) strains at various starting concentrations (based on OD_600_) with 0.1% v/v dimethyl sulfoxide (DMSO). We found that wells inoculated to a starting OD_600_ of 0.0075 resulted in excellent well-to-well consistency and reproducible growth after 24 h (**Fig. 1E**).

We then identified concentrations of phleomycin and herboxidiene that would achieve maximum growth inhibition under these conditions for use as positive controls. Using the 384-well plate format, we assayed concentration gradients of phleomycin (**Fig. 1F**; an anti-microbial that should prevent growth of all the yeast strains used in these studies) and herboxidiene (**Fig. 1G**; a splicing inhibitor that should prevent growth of the Hs-Hsh155 strain but not others). We found that robust growth inhibition could be obtained using concentrations of 16 µg/mL or 2 µM of phleomycin or herboxidiene, respectively.

### Pilot Screening of FDA-Approved Drugs

To evaluate this approach, we first screened the Selleck Chem FDA-approved drug library, which contains 1078 chemically diverse compounds. This screen was conducted in duplicate using the WT and Hs-Hsh155 yeast strains with the primary objective being to assess the sensitivity and specificity of yeast growth as a readout for identifying compounds of interest. As a basis for selecting potential growth inhibitors, we decided to choose those that reproducibly inhibited yeast growth in at least one strain by ≥ 30% relative to the corresponding DMSO controls.

Using this approach, we identified 18 compounds that inhibited yeast growth in at least one of the strains (**Fig. 1H, I**). Many of these are known antifungal drugs (*e.g.,* caspofungin, clotrimazole); so, their identification was expected. In addition to antifungals, we also identified members of other drug classes, including an antidepressant, immunosuppressants, a cognition-enhancing medication, and an antihistamine (**Supp. Table S1**). Importantly, the assay’s positive controls, phleomycin and herboxidiene were successfully identified, while the negative control, DMSO, was not. Herboxidiene was also identified only in the humanized, Hs-Hsh155 strain, not the WT-Hsh155 strain. These findings highlight the platform’s capability to not only identify compounds that inhibit yeast growth but also distinguish between compounds that selectively inhibit growth of one yeast strain over another via splicing modulation.

### Primary Screen Results

Based on results from the Selleck FDA-approved drug library, we expanded our screen to involve approximately 35,000 compounds sourced from four structurally diverse libraries: the RIKEN-Pilot Natural Products Depository (NPDepo), Life Chemicals 4 (LC4), and National Cancer Institute’s (NCI) Developmental Therapeutics Program (DTP) and Experimental Program (NExT) Diversity libraries. We also included the WT-Brr2 and Hs-Brr2 strains in the screen in addition to WT-Hsh155 and Hs-Hsh155. In these screens, we calculated a Z’ factor for each plate to assess screen reliability ^24^. Z’ values were calculated using responses from wells containing DMSO controls (no growth inhibition) vs. phleomycin (strong growth inhibition). All plate Z’ factor values exceed the cutoff of 0.5, indicating a high-quality screen (**Fig. 2A**).

**Figure 2.**
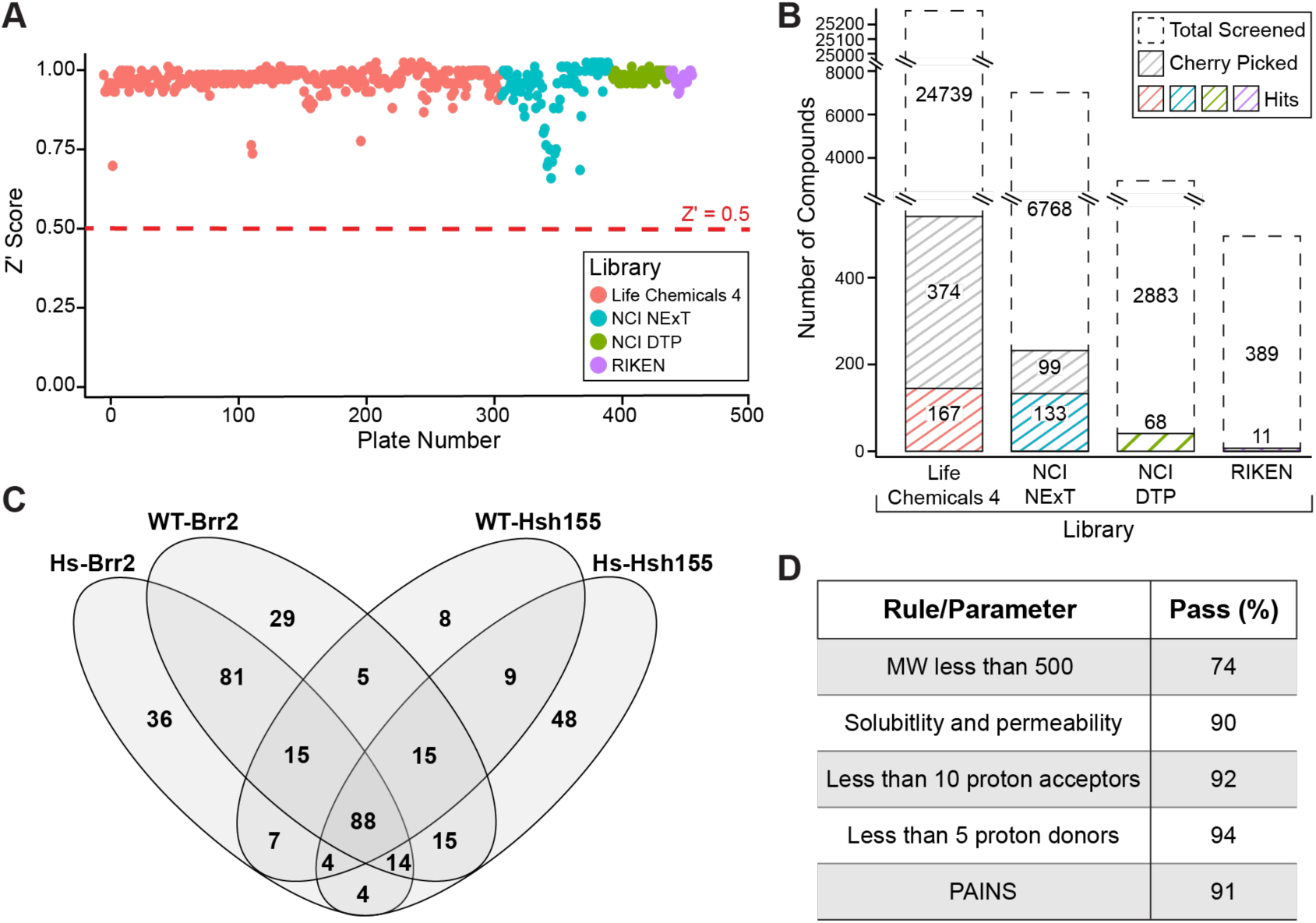
Primary Screen Results. **(A)** Scatterplot of Z’ scores for each screening plate. Red dotted line marks the quality threshold (Z’ = 0.5). Libraries are color-coded as indicated. **(B)** Stacked bar graph showing the number of compounds screened per library, categorized as total library compounds, those cherry-picked for subsequent replicates and rescreened but not identified as hits (grey diagonals), and primary hits (colored diagonals, which were derived from those that were cherry-picked for the Life Chemicals 4 and NCI NExT libraries). **(C)** Venn diagram illustrating overlap of primary hits across four yeast strains, highlighting shared and strain-specific compounds. **(D)** Table summarizing the percentage of primary hits that meet Lipinski’s Rule of Five and PAINS (Pan Assay Interference Compounds) filters.

Due to the sizes of the LC4 and NCI NExT libraries, we conducted an initial screen using all four strains and then selected compounds that showed ≥ 30% growth inhibition. We then screened these selected compounds a further two times, and those that consistently inhibited growth by at least 30% were designated as “hits” from the primary screens of these libraries. For the NCI DTP and RIKEN libraries, we screened the entire libraries three times in all four yeast strains. In total, 379 compounds were identified based on our metrics (**Fig. 2B**). We then separated the compounds based on which strain(s) growth inhibition was observed (**Fig. 2C**). While some compounds (88) appeared to have general growth inhibitory properties for yeast and inhibited growth of all strains, most of the compounds were selective for inhibiting growth in just one strain or a subset. This suggests that the chosen yeast strains were indeed sensitized to different compounds.

We further assessed the drug-likeness of these hits by investigating their structural and physicochemical properties. Using *in silico* cheminformatics analysis with SwissADME^33^, we clustered the lead candidates according to Lipinski’s rule of five parameters^34^ : molecular weight <500 g/mol, log*P* < 5 (where *P* is the partition coefficient in octanol vs. water), proton donors < 5, and proton acceptors < 10 (**Fig. 2D**). Approximately 70% of the hits demonstrated potential drug-likeness, possessing the chemical and physical properties conducive to oral bioavailability. We were unable to evaluate hits obtained from the Riken drug library due to lack of provided structural information. Finally, we analyzed the hits in terms of their potential presence as false positives by identifying any that are among the Pan Assay Interference Compounds (PAINS) class, which often appear as frequent hits in HTS assays^35^. The majority of our hits (91%) pass PAINS criteria, meaning they are not members of this class.

### Secondary Screen Optimization and Results

We next developed a secondary screen to discriminate between compounds in which growth inhibition is correlated with accumulation of unspliced pre-mRNAs and those which cause growth inhibition but do not generally inhibit splicing. We incorporated a fluorescent protein reporter gene (Venus) at the endogenous *URA3* locus in the WT-Hsh155, Hs-Hsh155, WT-Brr2, and Hs-Brr2 strains. We employed two different reporter gene constructs (**Fig. 3A**, **B**): SPLIF (splicing-in-frame) and SPLOOF (splicing-out-of-frame). The SPLIF reporter serves as a control, generating a Venus protein upon translation of the spliced mRNA. In contrast, when splicing is inhibited, Venus expression diminishes due to the inclusion of the intron and disruption of the reading frame (**Fig. 3A**). Conversely, the SPLOOF reporter only allows for the Venus protein production if the pre-mRNA is not spliced and the intron is retained (**Fig. 3B**). Thus, splicing modulators can be identified by observing decreases in fluorescence due to loss of Venus protein production in SPLIF-containing strains and increases in fluorescence due to production of Venus protein in SPLOOF-containing strains.

**Figure 3.**
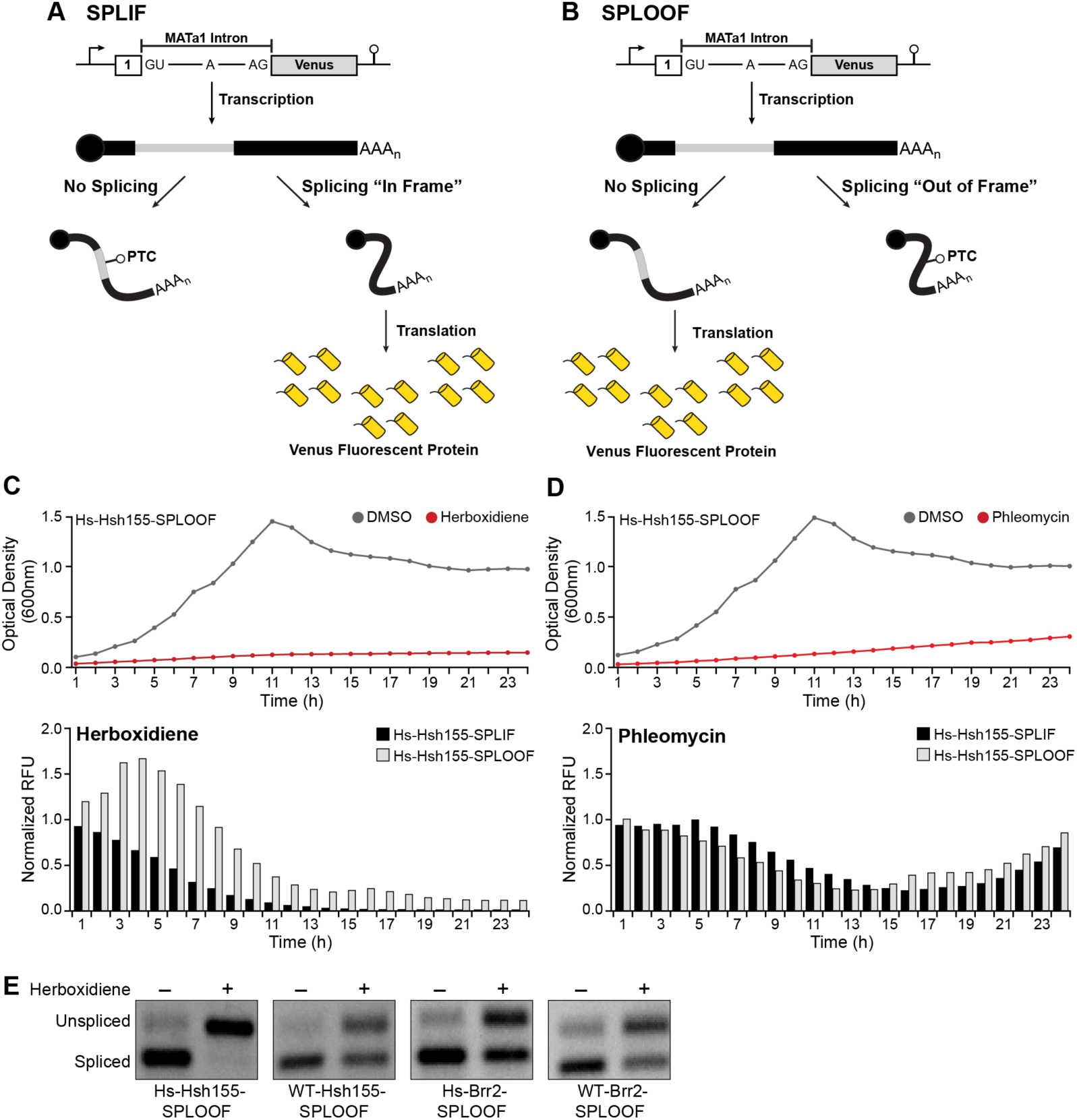
Secondary Screen Optimization. **(A, B)** Schematic of splicing inhibition reporters used in secondary screening. Both constructs include the yeast MATa1 intron inserted into the ORF for the Venus fluorescent protein gene. In the SPLIF reporter, proper splicing produces high Venus fluorescence, which decreases with splicing inhibition. In the SPLOOF reporter, splicing disrupts the reading frame, suppressing Venus expression; splicing inhibition restores in-frame expression, increasing fluorescence. **(C, D)** (Top) Optical density measurement time courses (OD_600_) of the Hs-Hsh155-SPLOOF strain upon treatment with either DMSO (grey) or herboxidiene or phleomycin (red). (Bottom) Fluorescence measurement time courses of Hs-Hsh155-SPLIF (black bars) or Hs-Hsh155-SPLOOF (grey bars) following treatment with 2 µM herboxidiene or 16 µg/mL phleomycin, normalized to DDMSO controls. **(E)** RT-PCR validation of splicing inhibition in SPLOOF strains treated with DMSO or 2 µM herboxidiene, confirming retention of the MATa1 first intron (unspliced band) upon addition of drug.

To validate this secondary screen, we measured changes in both cell culture fluorescence (normalized to the DMSO control) and optical density for 24 h for Hs-Hsh155 strains containing SPLIF or SPLOOF reporters in the presence of DMSO, phleomycin, or herboxidiene (**Fig. 3C**, **D**). As expected, we observed significant growth inhibition, as measured by the OD_600_, for these strains in the presence of phleomycin (16 µg/mL) or herboxidiene (2 µM) but not in the presence of DMSO (shown in **Fig. 3C** Hs-Hsh155-SPLOOF). In the presence of herboxidiene, we observed a transient increase in fluorescence lasting from ∼1-7h during drug treatment for the strain carrying the SPLOOF reporter and a corresponding decrease in fluorescence from the strain with the SPLIF reporter. In contrast, we only observed decreases in fluorescence when cells were exposed to phleomycin. These results show that addition of a known splicing modulator can cause changes in culture fluorescence using the SPLIF/SPLOOF reporter system.

To verify that cellular pre-mRNA was indeed accumulating upon treatment with herboxidiene, we used RT-PCR to determine relative abundances of unspliced and spliced junctions for the first intron of endogenous MATa1 transcript (**Fig. 3E**). These results confirmed that herboxidiene was causing unspliced transcript accumulation in the Hsh155- and Brr2-modified yeast strains. Interestingly, we could also observe some accumulation of the unspliced RNA in strains not expected to be sensitive to herboxidiene (WT-Hsh155 and the Brr2-modified strains), albeit to a lower level than the sensitized strain (Hs-Hsh155). This suggests that herboxidiene can at least partially inhibit yeast pre-mRNA splicing under these conditions, even when little-to-no growth inhibition is observed. This is consistent with results recently reported by Hunter and colleagues^9^.

We then used this fluorescence-based screen to assay all 379 compounds identified in the primary screen in eight different yeast strains (Hsh155- and Brr2-modified strains, each with either the SPLIF or SPLOOF reporter) in duplicate. In each case, yeast were exposed to the compounds for a total of 24 h. We used a threshold for splicing modulation as the presence of a normalized (relative to DMSO) fluorescence intensity signal > 1.0 for stains with the SPLOOF reporter along with a signal of < 1.0 for the corresponding condition with the SPLIF reporter. In addition, we selected for compounds that caused changes in fluorescence that lasted at least for ten consecutive cycles of measurement, or ∼2.5h. From the starting set of 379 compounds, we identified 11 that met all secondary screening criteria (**Supp. Table S2**).

When analyzing these compounds, we noted that the RIKEN library was the only one that did not yield any secondary hits (**Fig. 4A**). Hits were evenly distributed among the other libraries with no particular library more likely to contain a hit than any other based only on the distribution of the primary screen hits. In terms of total library compounds, the greatest hit rates were obtained from the NCI NeXT and DTP libraries (*cf.* **Fig. 2B**, **Fig. 4A**). Among the 11 hits, one structural scaffold was represented by multiple hits: adenosine analogs. Both 3’-deoxyadenosine (cordycepin) and 9-β-D-erythrofuranosyladenine (EFA), resulted in changes in splicing reporter fluorescence in all of the tested strains (**Fig. 4B**, **C**; **Supp. Table S2**). Additional scaffolds included benzothiazoles and benzoxazoles, which showed some strain specificity in the Hs-Brr2 (oxazoles) and WT-Hsh155 (thiazole) backgrounds. Other hits were structurally unique, and their strain specificities are detailed in Supplementary Table S2.

**Figure 4.**
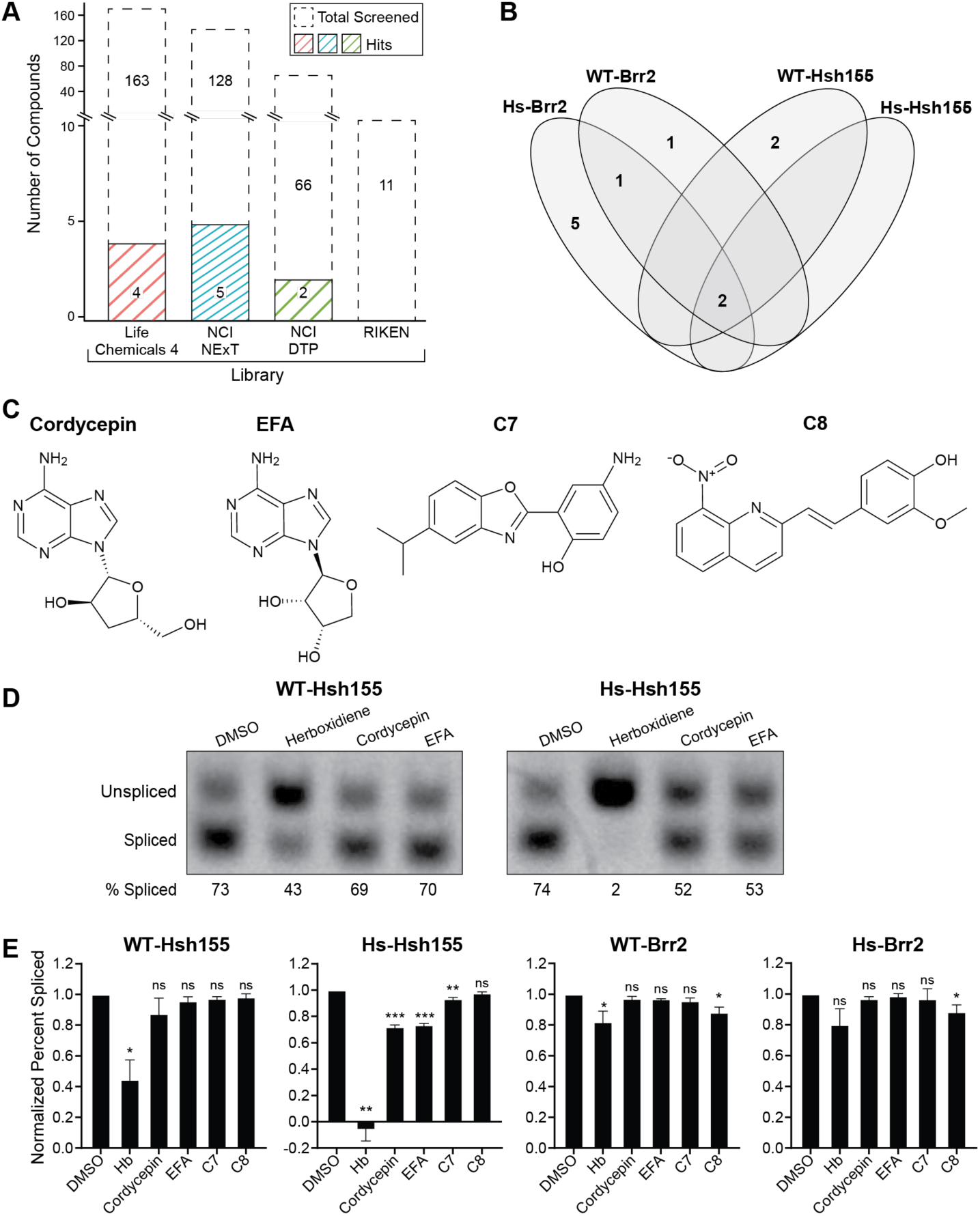
Secondary Screening Results. **(A)** Stacked bar graph showing the number of compounds screened per library using the secondary screen and the corresponding number of hits. **(B)** Venn diagram showing overlap of secondary screen hits between strains. **(C)** Chemical structures of cordycepin, EFA, compound C7, and compound C8. **(D)** Representative RT-PCR results and quantification of the splicing of the endogenous MATa1 gene first intron in each strain treated with 10 µM of each compound. Values are normalized to DMSO. Error bars represent standard deviation (*N*= 3). Statistical significance of the effects of compound treatment vs. the DMSO controls was calculated using an unpaired two-tailed Welch’s *t*-test (\**p* < 0.05, \*\**p* < 0.005, \*\*\**p* < 0.001).

### Additional Validation and Characterization of Secondary Screen Hits

To further confirm the splicing modulatory potential of the 11 hits, we assessed removal of the first intron of the endogenous MATa1 transcript by RT-PCR for each compound in the WT-Hsh155, Hs-Hsh155, WT-Brr2, and Hs-Brr2 strains. We compared the effects of each compound to herboxidiene and DMSO controls. Four of the 11 compounds that passed secondary screening demonstrated significant splicing inhibition of the endogenous MATa1 transcript (**Fig. 4C**, **D**). Cordycepin, EFA, and compound C7 caused the most substantial accumulation of unspliced RNA in the Hs-Hsh155 strain, while compound C8 reduced splicing efficiency in WT-Brr2 and Hs-Brr2 strains. It is possible that the remaining 7 compounds also inhibit splicing of other RNAs besides that in the MATa1 transcript tested here; however, we did not explore this further. Nonetheless, we conclude that our screen was successfully able to identify yeast splicing modulators that function *in vivo* and that the efficacies of these modulators are strain-dependent.

### Splicing Factor Mutant Cancer Cells are Sensitive to Compound C7

Finally, we decided to test if any of the identified compounds also exhibited splicing modulatory activity in human cells. Previous studies have demonstrated that cancer cells harboring mutant splicing factors are more susceptible to splicing modulation than their WT counterparts^36^. Inspired by this, we tested four compounds (those shown in **Fig. 4C**) for antiproliferative activity against isogenic K562 cancer cells expressing either wildtype SRSF2 or the hematologic malignancy-associated *SRSF2^P95H^* mutation^26^. Cells were treated with increasing concentrations of each drug, and relative cell viability was assessed after seven days. We did not find any significant differences in antiproliferative activity for cordycepin, EFA, or compound C8 between the SRSF2^WT^ and SRSF2^P95H^ cells (data not shown). However, we observed the SRSF2^P95H^ cells exhibited reduced proliferation relative to SRSF2^WT^ cells in response to compound C7 (**Fig. 5A**, **B**). We note that compound C7 has been reported as a potential luciferase inhibitor^37^, which could interfere with the proliferation assay. However, we did not detect any inhibition of luciferase activity in control experiments using the Cell-TiterGlo kit (data not shown).

**Figure 5.**
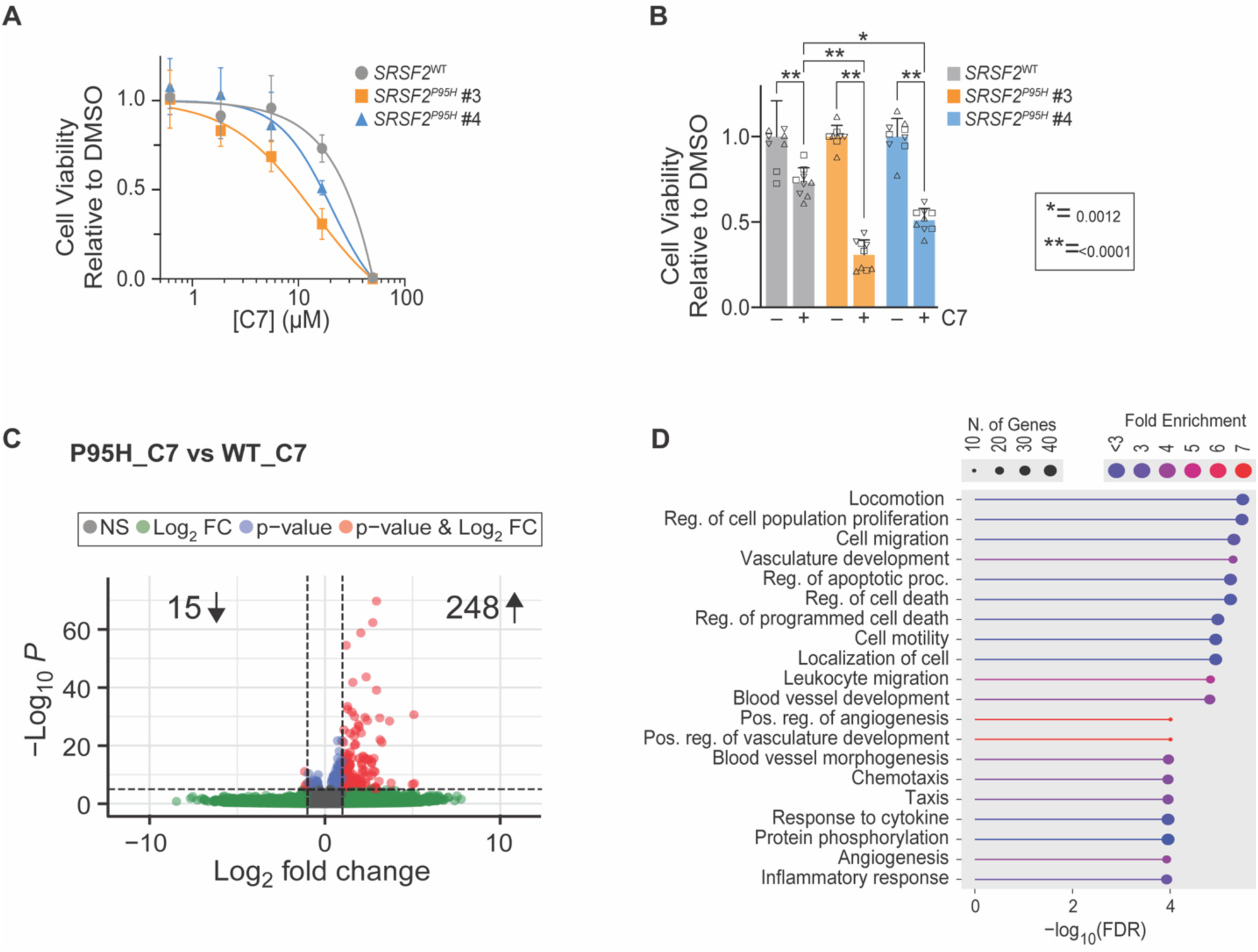
Splicing Mutant Cancer Cells are Sensitized to Compound C7. **(A)** Relative viability of K562 SRSF2^WT/WT^ and SRSF2^WT/P95H^ K562 cells in tissue culture treated with increasing concentrations of C7 for 7 days. Data points represent the mean of 3 biological replicates and error bars represent standard deviation. Two independently-obtained clones of the SRSF2^WT/P95H^ cell line were tested (clones #3 and #4). (B) Relative survival of K562 SRSF2^WT^ and SRSF2^P95H^ treated with 16.7 µM of C7 for 7 days. Bars represent the mean of 9 technical replicates across 3 biological replicates and error bars represent standard deviation. Two-way ANOVA with Sidak’s multiple comparison test. **(C)** Volcano plot displaying uniquely down (left side) and up-regulated DEGs (right side) in K562 SRSF2 ^WT/P95H^ cells vs. WT when treated with C7 relative to DMSO controls for each. DEGs were defined as those with log₂ fold change < -1 or >1 and a -log₁₀ *p*-value >0.05 (red dots). **(D)** Gene ontology (GO) enrichment analysis of the upregulated DEGs.

### Compound C7 Does Not Generally Inhibit pre-mRNA Splicing *in vitro* but Changes Gene Expression *in vivo*

To further explore the mechanism by which SRSF2-mutant cells are sensitive to compound C7, we analyzed pre-mRNA splicing *in vitro* and *in vivo*. We were unable to observe any significant reduction in splicing of the well-characterized AdML pre-mRNA substrate in human HeLa cell nuclear extract (a standard extract and substrate for *in vitro* assays of the human splicing machinery) at concentrations up to 200 µM of compound C7 (**Supp. Fig. S1**). This suggests that either compound C7 does not target the core splicing machinery directly or that any impact on splicing may be cell type or transcript-specific and not able to be reconstituted *in vitro* using this extract and/or substrate.

To assess transcriptome changes caused by compound C7 in K562 cells, we carried out RNA sequencing on poly-A selected RNA isolated from SRSF2 WT or P95H mutant cells after exposure to DMSO or C7 (16.7 µM for 24h). We first analyzed the results in terms of differentially expressed genes (DEGs), which we defined as those with a log_2_-fold change of < -1 (downregulated) or > 1 (upregulated) and an adjusted *p*-value < 0.05.

As expected, the most prominent changes in DEG occurred in comparison of the SRSF2 WT and P95H mutant cells even in the absence of drug treatment (**Supp. Fig. S2**). This is consistent with previous findings that the SRSF2^P95H^ mutation by itself alters gene expression and splicing^38^.

When comparing each strain’s response to C7, we found that C7 caused relatively few uniquely downregulated DEGs in the SRSF2^P95H^ mutant cells relative to SRSF2^WT^ (15 DEGs, **Fig. 5C**). However, it resulted in many more upregulated genes in the SRSF2^WT/P95H^ cells including those involved in regulation of apoptosis and cell death (248 DEGs; **Fig. 5C**, **D**). It is possible that dysregulation of this process leads to the decrease in proliferation observed in SRSF2^WT/P95H^ cells relative to WT upon treatment with C7.

### Changes in pre-mRNA Splicing due to Treatment with Compound C7

We then looked at the changes in splicing due to compound C7 using rMATS to perform differential splicing analysis^28^. Again, consistent with prior studies, we observed many changes in splicing due to the SRSF2^P95H^ mutation itself, even in the absence of compound treatment (**Sup. Fig. S3**)^38^. Therefore, we focused on the events in each cell line induced by compound C7 relative to the DMSO control **(Fig. 6A**). The most common event types were skipped exons; however, we also observed changes in retained introns, mutually exclusive exons, and splice site usage. Compound C7 induced more changes in alternative splicing outcomes in the SRSF2^P95H^ mutant cells relative to those with SRSF2^WT^.

**Figure 6.**
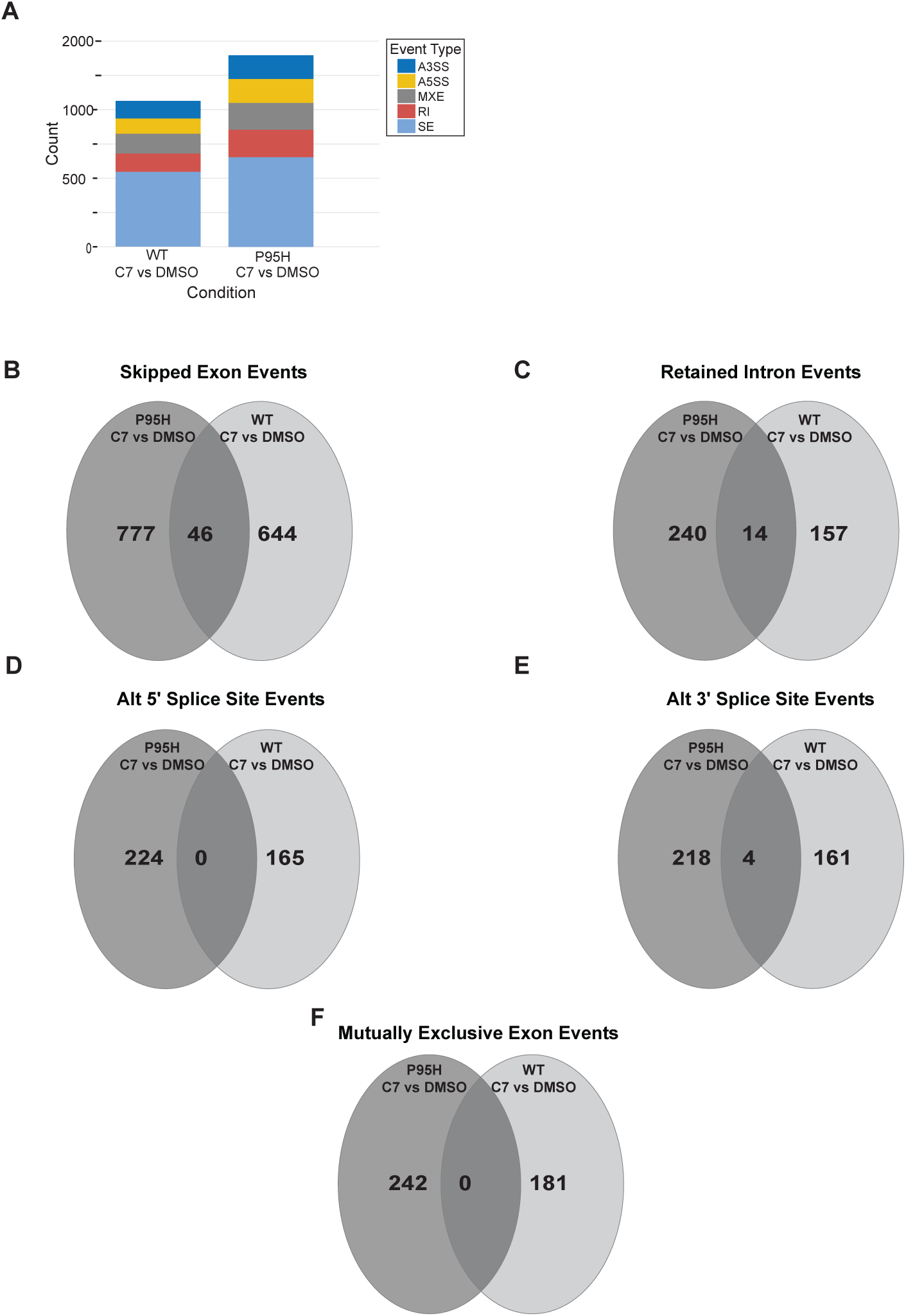
C7 Treatment Induces Changes in pre-mRNA Splicing. **(A)** Stacked bar graph showing the distribution of novel alternative splicing events found in WT or P95H cells upon treatment with C7 relative to the corresponding DMSO control. A3SS/A5SS: alternative 3’/5’ splice sites; MXE: mutually exclusive exons; RI: retained introns; SE: skipped exons. **(B–F)** Venn diagrams showing unique splicing events in the P95H mutant cells (dark grey) in comparison to WT (light grey) upon treatment with C7 relative to the DMSO control for each. Overlapping events were common to both cell lines upon compound treatment.

We created Venn diagrams to quantify the numbers of unique or shared events in each cell line due to compound treatment (**Fig. 6B-F**). This analysis revealed that the majority of the changes in alternative splicing due to compound C7 were unique to each cell type and relatively few events were shared. GO-term analysis of the genes with alternative splicing events unique to the SRSF2 P95H mutant cells upon C7 treatment relative to WT showed that the splicing changes impacted a wide range of processes especially those involving gene expression including transcriptional regulation, DNA and RNA metabolism, and translation (**Supp. Table S3**). As with upregulation of genes involved in apoptosis, dysregulation of mRNA isoform generation could lead to the observed proliferative defects in the SRSF2^P95H^ mutant cells. In summary, we conclude that compound C7 modulates splicing in SRSF2^WT^ and SRSF2^P95H^ mutant cells differently, it perturbs alternative splicing regulation of a greater number of RNAs in SRSF2^P95H^ cells, and that these transcriptome-wide impacts that may result in impaired cellular proliferation.

## CONCLUSION

In this work, we created a yeast cell-based high-throughput screening platform that identified four *in vivo* splicing modulators. These compounds inhibit growth of *S. cerevisiae* and can cause accumulation of unspliced MATa1 transcripts in yeast. Further analysis of these compounds demonstrated that one, compound C7, reduces proliferation of K562 cells harboring the cancer-associated SRSF2^P95H^ mutation and that these splicing factor-mutant cells are more sensitive to this compound than their wildtype counterparts. The increased sensitivity could be due to changes in regulation of genes involved in processes such as apoptosis and/or reprogramming of alternative splicing.

A significant advantage of our screening platform is its simplicity, cost-effectiveness, and scalability. Unlike mammalian cell-based assays, our yeast system can be easily set-up on a lab bench without needing a biosafety cabinet. Yeasts grow much more rapidly than mammalian cells and in much less expensive growth media. This allows for facile, low-cost, and rapid identification of splicing modulators. A limitation of our studies is that we have not identified the precise targets of these compounds in yeast or human cells. It is possible that these compounds may exert their effects via indirect mechanisms. For example, by changing transcription, chromatin structure, or RNA export. It is interesting to note that two of the identified compounds, cordycepin and EFA, are adenosine analogs. We speculate that these could inhibit splicing by associating with nucleotide binding pockets within the splicing machinery (such as the branch site adenosine pocket on Hsh155/SF3B1)^39^ by interfering with ATP binding to the highly conserved splicesomal ATPases ^40^ or by indirectly impacting splicing through changes in coupled RNA processing events such as transcription or polyadenylation. Future work could focus on target deconvolution for these molecules as well as expanding studies of C7 for potential therapeutic applications.

## Acknowledgments

SLL, HV, and AAH were funded by a EvansMDS Discovery research grant from the Edward P. Evans Foundation. SLL was supported by the Genetics Training Program (NIH 5T32GM007133) and the SciMed Graduate Research Scholars Fellowship. YCC was supported by the Targets of Cancer Training Program (NIH T32CA009138). MSJ was supported by the National Institutes of Health (R01GM72649). HDN is supported by grants from the Masonic Cancer Center, Edward P. Evans Foundation, American Society of Hematology, the NIH’s National Heart, Lung, and Blood Institute (R01HL163011), and the 2022 AACR Career Development Award to Further Diversity, Equity, and Inclusion in Cancer Research, which is supported by Merck, grant number 22–20–68-NGUY. We thank the UW-Madison small molecule screening facility for instrument access. We also thank Dr. Manny Ares for splicing reporter plasmids. Finally, we thank Laura Vanderploeg for figure editing.

## References

(1) Dvinge, H.; Kim, E.; Abdel-Wahab, O.; Bradley, R. K. RNA splicing factors as oncoproteins and tumour suppressors. Nat Rev Cancer 2016, 16 (7), 413–430. DOI: 10.1038/nrc.2016.51 From NLM Medline. Love, S. L.; Emerson, J. D.; Koide, K.; Hoskins, A. A. Pre-mRNA splicing-associated diseases and therapies. RNA Biol 2023, 20 (1), 525–538. DOI: 10.1080/15476286.2023.2239601 From NLM Medline.

(2) Effenberger, K. A.; Urabe, V. K.; Jurica, M. S. Modulating splicing with small molecular inhibitors of the spliceosome. Wiley Interdiscip Rev RNA 2017, 8 (2), 19. DOI: 10.1002/wrna.1381 From NLM Medline.

(3) Dmitriev, S. E.; Vladimirov, D. O.; Lashkevich, K. A. A Quick Guide to Small-Molecule Inhibitors of Eukaryotic Protein Synthesis. Biochemistry (Mosc) 2020, 85 (11), 1389–1421. DOI: 10.1134/S0006297920110097 From NLM Medline.

(4) Ji, T.; Yang, Y.; Yu, J.; Yin, H.; Chu, X.; Yang, P.; Xu, L.; Wang, X.; Hu, S.; Li, Y.;, et al. Targeting RBM39 through indisulam induced mis-splicing of mRNA to exert anti-cancer effects in T-cell acute lymphoblastic leukemia. J Exp Clin Cancer Res 2024, 43 (1), 205. DOI: 10.1186/s13046-024-03130-8 From NLM Medline. Singh, S.; Quarni, W.; Goralski, M.; Wan, S.; Jin, H.; Van de Velde, L. A.; Fang, J.; Wu, Q.; Abu-Zaid, A.; Wang, T.; et al. Targeting the spliceosome through RBM39 degradation results in exceptional responses in high-risk neuroblastoma models. Sci Adv 2021, 7 (47), eabj5405. DOI: 10.1126/sciadv.abj5405 From NLM PubMed-not-MEDLINE. Xu, Y.; Nijhuis, A.; Keun, H. C. RNA-binding motif protein 39 (RBM39): An emerging cancer target. Br J Pharmacol 2022, 179 (12), 2795–2812. DOI: 10.1111/bph.15331 From NLM Medline.

(5) Cretu, C.; Agrawal, A. A.; Cook, A.; Will, C. L.; Fekkes, P.; Smith, P. G.; Luhrmann, R.; Larsen, N.; Buonamici, S.; Pena, V. Structural Basis of Splicing Modulation by Antitumor Macrolide Compounds. Mol Cell 2018, 70 (2), 265–273 e268. DOI: 10.1016/j.molcel.2018.03.011 From NLM Medline.

(6) Cretu, C.; Gee, P.; Liu, X.; Agrawal, A.; Nguyen, T. V.; Ghosh, A. K.; Cook, A.; Jurica, M.; Larsen, N. A.; Pena, V. Structural basis of intron selection by U2 snRNP in the presence of covalent inhibitors. Nat Commun 2021, 12 (1), 4491. DOI: 10.1038/s41467-021-24741-1 From NLM Medline.

(7) Carrocci, T. J.; Paulson, J. C.; Hoskins, A. A. Functional analysis of Hsh155/SF3b1 interactions with the U2 snRNA/branch site duplex. RNA 2018, 24 (8), 1028–1040. DOI: 10.1261/rna.065664.118 From NLM Medline.

(8) Hansen, S. R.; Nikolai, B. J.; Spreacker, P. J.; Carrocci, T. J.; Hoskins, A. A. Chemical Inhibition of Pre-mRNA Splicing in Living Saccharomyces cerevisiae. Cell Chem Biol 2019, 26 (3), 443–448 e443. DOI: 10.1016/j.chembiol.2018.11.008 From NLM Medline.

(9) Hunter, O.; Talkish, J.; Quick-Cleveland, J.; Igel, H.; Tan, A.; Kuersten, S.; Katzman, S.; Donohue, J. P.; M. S. J.; Ares, M., Jr. Broad variation in response of individual introns to splicing inhibitors in a humanized yeast strain. RNA 2024, 30 (2), 149–170, Article. DOI: 10.1261/rna.079866.123 From NLM Medline.

(10) Rangan, R.; Huang, R.; Hunter, O.; Pham, P.; Ares, M.; Das, R. RNA structure landscape of *S. cerevisiae* introns. bioRxiv 2024, 2022.2007.2022.501175. DOI: 10.1101/2022.07.22.501175.

(11) Lazear, M. R.; Remsberg, J. R.; Jaeger, M. G.; Rothamel, K.; Her, H. L.; DeMeester, K. E.; Njomen, E.; Hogg, S. J.; Rahman, J.; Whitby, L. R.;, et al. Proteomic discovery of chemical probes that perturb protein complexes in human cells. Mol Cell 2023, 83 (10), 1725–1742 e1712. DOI: 10.1016/j.molcel.2023.03.026 From NLM Medline. Tellier, M.; Ansa, G.; Murphy, S. Isoginkgetin and Madrasin are poor splicing inhibitors. PLoS One 2024, 19 (10), e0310519. DOI: 10.1371/journal.pone.0310519 From NLM Medline. Sidarovich, A.; Will, C. L.; Anokhina, M. M.; Ceballos, J.; Sievers, S.; Agafonov, D. E.; Samatov, T.; Bao, P.; Kastner, B.; Urlaub, H.; et al. Identification of a small molecule inhibitor that stalls splicing at an early step of spliceosome activation. Elife 2017, 6. DOI: 10.7554/eLife.23533 From NLM Medline.

(12) Fuentes-Fayos, A. C.; Pérez-Gómez, J. M.; G-García, M. E.; Jiménez-Vacas, J. M.; Blanco-Acevedo, C.; Sánchez-Sánchez, R.; Solivera, J.; Breunig, J. J.; Gahete, M. D.; Castaño, J. P.;, et al. SF3B1 inhibition disrupts malignancy and prolongs survival in glioblastoma patients through BCL2L1 splicing and mTOR/ß-catenin pathways imbalances. Journal of Experimental & Clinical Cancer Research 2022 41:1 2022, 41 (1). DOI: 10.1186/s13046-022-02241-4. Lopez-Oreja, I.; Gohr, A.; Playa-Albinyana, H.; Giro, A.; Arenas, F.; Higashi, M.; Tripathi, R.; Lopez-Guerra, M.; Irimia, M.; Aymerich, M.; et al. SF3B1 mutation-mediated sensitization to H3B-8800 splicing inhibitor in chronic lymphocytic leukemia. Life Sci Alliance 2023, 6 (11), 25. DOI: 10.26508/lsa.202301955 From NLM Medline. Maji, D.; Grossfield, A.; Kielkopf, C. L. Structures of SF3b1 reveal a dynamic Achilles heel of spliceosome assembly: Implications for cancer-associated abnormalities and drug discovery. Biochim Biophys Acta Gene Regul Mech 2019, 1862 (11-12), 194440. DOI: 10.1016/j.bbagrm.2019.194440 From NLM Medline.

(13) Seiler, M.; Yoshimi, A.; Darman, R.; Chan, B.; Keaney, G.; Thomas, M.; Agrawal, A. A.; Caleb, B.; Csibi, A.; Sean, E.;, et al. H3B-8800, an orally available small-molecule splicing modulator, induces lethality in spliceosome-mutant cancers. Nat Med 2018, 24 (4), 497–504. DOI: 10.1038/nm.4493 From NLM Medline.

(14) Bowling, E. A.; Wang, J. H.; Gong, F.; Wu, W.; Neill, N. J.; Kim, I. S.; Tyagi, S.; Orellana, M.; Kurley, S. J.; Dominguez-Vidana, R.;, et al. Spliceosome-targeted therapies trigger an antiviral immune response in triple-negative breast cancer. Cell 2021, 184 (2), 384–403 e321. DOI: 10.1016/j.cell.2020.12.031 From NLM Medline. Agrawal, A. A.; Yu, L.; Smith, P. G.; Buonamici, S. Targeting splicing abnormalities in cancer. Curr Opin Genet Dev 2018, 48, 67–74. DOI: 10.1016/j.gde.2017.10.010 From NLM Medline.

(15) Dart, A. Targeting aberrant splicing. Nat Rev Cancer 2023, 23 (10), 653, Editorial Material; Early Access. DOI: 10.1038/s41568-023-00620-3 From NLM PubMed-not-MEDLINE.

(16) Wheeler, E. C.; Martin, B. J. E.; Doyle, W. C.; Neaher, S.; Conway, C. A.; Pitton, C. N.; Gorelov, R. A.; Donahue, M.; Jann, J. C.; Abdel-Wahab, O.;, et al. Splicing modulators impair DNA damage response and induce killing of cohesin-mutant MDS and AML. Sci Transl Med 2024, 16 (728), eade2774. DOI: 10.1126/scitranslmed.ade2774 From NLM Medline. Yamauchi, H.; Nishimura, K.; Yoshimi, A. Aberrant RNA splicing and therapeutic opportunities in cancers. Cancer Sci 2022, 113 (2), 373–381. DOI: 10.1111/cas.15213 From NLM Medline.

(17) Lu, S. X.; De Neef, E.; Thomas, J. D.; Sabio, E.; Rousseau, B.; Gigoux, M.; Knorr, D. A.; Greenbaum, B.; Elhanati, Y.; Hogg, S. J.;, et al. Pharmacologic modulation of RNA splicing enhances anti-tumor immunity. Cell 2021, 184 (15), 4032–4047 e4031. DOI: 10.1016/j.cell.2021.05.038 From NLM Medline. Kim, W. J.; Crosse, E. I.; De Neef, E.; Etxeberria, I.; Sabio, E. Y.; Wang, E.; Bewersdorf, J. P.; Lin, K. T.; Lu, S. X.; Belleville, A.; et al. Mis-splicing-derived neoantigens and cognate TCRs in splicing factor mutant leukemias. Cell 2025. DOI: 10.1016/j.cell.2025.03.047 From NLM Publisher.

(18) Owens, D. D. G.; Caulder, A.; Frontera, V.; Harman, J. R.; Allan, A. J.; Bucakci, A.; Greder, L.; Codner, G. F.; Hublitz, P.; McHugh, P. J.;, et al. Microhomologies are prevalent at Cas9-induced larger deletions. Nucleic Acids Res 2019, 47 (14), 7402–7417. DOI: 10.1093/nar/gkz459 From NLM Medline.

(19) Goldstein, A. L.; McCusker, J. H. Three new dominant drug resistance cassettes for gene disruption in Saccharomyces cerevisiae. Yeast 1999, 15 (14), 1541–1553. DOI: 10.1002/(SICI)1097-0061(199910)15:14<1541::AID-YEA476>3.0.CO;2-K From NLM Medline.

(20) Lee, M. E.; DeLoache, W. C.; Cervantes, B.; Dueber, J. E. A Highly Characterized Yeast Toolkit for Modular, Multipart Assembly. ACS Synth Biol 2015, 4 (9), 975–986. DOI: 10.1021/sb500366v From NLM Medline.

(21) Amberg, D. C.; Burke, D. J. Classical Genetics with Saccharomyces cerevisiae. Cold Spring Harb Protoc 2016, 2016 (5). DOI: 10.1101/pdb.top077628 From NLM Medline.

(22) Cheng, T. H.; Chang, C. R.; Joy, P.; Yablok, S.; Gartenberg, M. R. Controlling gene expression in yeast by inducible site-specific recombination. Nucleic Acids Res 2000, 28 (24), E108. DOI: 10.1093/nar/28.24.e108 From NLM Medline.

(23) Bland, J. M.; Altman, D. G. Statistical methods for assessing agreement between two methods of clinical measurement. Lancet 1986, 1 (8476), 307–310. From NLM Medline.

(24) Zhang, J. H.; Chung, T. D.; Oldenburg, K. R. A Simple Statistical Parameter for Use in Evaluation and Validation of High Throughput Screening Assays. J Biomol Screen 1999, 4 (2), 67–73. DOI: 10.1177/108705719900400206 From NLM Publisher.

(25) Effenberger, K. A.; Perriman, R. J.; Bray, W. M.; Lokey, R. S.; Ares, M., Jr.; Jurica, M. S. A high-throughput splicing assay identifies new classes of inhibitors of human and yeast spliceosomes. J Biomol Screen 2013, 18 (9), 1110–1120. DOI: 10.1177/1087057113493117 From NLM Medline.

(26) Liu, Z. S.; Sinha, S.; Bannister, M.; Song, A.; Arriaga-Gomez, E.; McKeeken, A. J.; Bonner, E. A.; Hanson, B. K.; Sarchi, M.; Takashima, K.;, et al. R-Loop Accumulation in Spliceosome Mutant Leukemias Confers Sensitivity to PARP1 Inhibition by Triggering Transcription-Replication Conflicts. Cancer Res 2024, 84 (4), 577–597. DOI: 10.1158/0008-5472.CAN-23-3239 From NLM Medline.

(27) Love, M. I.; Huber, W.; Anders, S. Moderated estimation of fold change and dispersion for RNA-seq data with DESeq2. Genome Biology 2014, 15 (12). DOI: 10.1186/s13059-014-0550-8.

(28) Shen, S.; Park, J. W.; Lu, Z. X.; Lin, L.; Henry, M. D.; Wu, Y. N.; Zhou, Q.; Xing, Y. rMATS: robust and flexible detection of differential alternative splicing from replicate RNA-Seq data. Proc Natl Acad Sci U S A 2014, 111 (51), E5593–5601. DOI: 10.1073/pnas.1419161111 From NLM Medline.

(29) Ge, S. X.; Jung, D.; Yao, R. ShinyGO: a graphical gene-set enrichment tool for animals and plants. Bioinformatics 2020, 36 (8), 2628–2629. DOI: 10.1093/bioinformatics/btz931 From NLM Medline.

(30) Iwatani-Yoshihara, M.; Ito, M.; Klein, M. G.; Yamamoto, T.; Yonemori, K.; Tanaka, T.; Miwa, M.; Morishita, D.; Endo, S.; Tjhen, R.;, et al. Discovery of Allosteric Inhibitors Targeting the Spliceosomal RNA Helicase Brr2. J Med Chem 2017, 60 (13), 5759–5771. DOI: 10.1021/acs.jmedchem.7b00461 From NLM Medline.

(31) Ito, M.; Iwatani, M.; Yamamoto, T.; Tanaka, T.; Kawamoto, T.; Morishita, D.; Nakanishi, A.; Maezaki, H. Discovery of spiro[indole-3,2’-pyrrolidin]-2(1H)-one based inhibitors targeting Brr2, a core component of the U5 snRNP. Bioorg Med Chem 2017, 25 (17), 4753–4767, Article. DOI: 10.1016/j.bmc.2017.07.017 From NLM Medline. Effenberger, K. A.; Urabe, V. K.; Prichard, B. E.; Ghosh, A. K.; Jurica, M. S. Interchangeable SF3B1 inhibitors interfere with pre-mRNA splicing at multiple stages. RNA 2016, 22 (3), 350–359. DOI: 10.1261/rna.053108.115 From NLM Medline.

(32) McMurray, M. A.; Thorner, J. Septin stability and recycling during dynamic structural transitions in cell division and development. Curr Biol 2008, 18 (16), 1203–1208. DOI: 10.1016/j.cub.2008.07.020 From NLM Medline.

(33) Daina, A.; Michielin, O.; Zoete, V. SwissADME: a free web tool to evaluate pharmacokinetics, drug-likeness and medicinal chemistry friendliness of small molecules. Sci Rep 2017, 7, 42717. DOI: 10.1038/srep42717 From NLM Medline.

(34) Lipinski, C. A.; Lombardo, F.; Dominy, B. W.; Feeney, P. J. Experimental and computational approaches to estimate solubility and permeability in drug discovery and development settings. Adv Drug Deliv Rev 2001, 46 (1-3), 3–26. DOI: 10.1016/s0169-409x(00)00129-0 From NLM Medline. Lipinski, C. A. Drug-like properties and the causes of poor solubility and poor permeability. J Pharmacol Toxicol Methods 2000, 44 (1), 235–249. DOI: 10.1016/s1056-8719(00)00107-6 From NLM Medline.

(35) Baell, J. B.; Holloway, G. A. New substructure filters for removal of pan assay interference compounds (PAINS) from screening libraries and for their exclusion in bioassays. J Med Chem 2010, 53 (7), 2719–2740. DOI: 10.1021/jm901137j From NLM Medline.

(36) Pellagatti, A.; Boultwood, J. Splicing factor mutations in the myelodysplastic syndromes: Role of key aberrantly spliced genes in disease pathophysiology and treatment. Adv Biol Regul 2023, 87, 100920. DOI: 10.1016/j.jbior.2022.100920 From NLM Medline. Lee, S. C.; Dvinge, H.; Kim, E.; Cho, H.; Micol, J. B.; Chung, Y. R.; Durham, B. H.; Yoshimi, A.; Kim, Y. J.; Thomas, M.; et al. Modulation of splicing catalysis for therapeutic targeting of leukemia with mutations in genes encoding spliceosomal proteins. Nat Med 2016, 22 (6), 672–678. DOI: 10.1038/nm.4097 From NLM Medline.

(37) PubChem Compound Summary for CID 833827, 4-Amino-2-[5-(propan-2-yl)-1,3-benzoxazol-2-yl]phenol. https://pubchem.ncbi.nlm.nih.gov/compound/833827 (accessed.

(38) Jia, W.; Guo, X.; Wei, Y.; Liu, J.; Can, C.; Wang, R.; Yang, X.; Ji, C.; Ma, D. Clinical and prognostic profile of SRSF2 and related spliceosome mutations in patients with acute myeloid leukemia. Mol Biol Rep 2023, 50 (8), 6601–6610. DOI: 10.1007/s11033-023-08597-w From NLM Medline. Kim, E.; Ilagan, J. O.; Liang, Y.; Daubner, G. M.; Lee, S. C.; Ramakrishnan, A.; Li, Y.; Chung, Y. R.; Micol, J. B.; Murphy, M. E.; et al. SRSF2 Mutations Contribute to Myelodysplasia by Mutant-Specific Effects on Exon Recognition. Cancer Cell 2015, 27 (5), 617–630. DOI: 10.1016/j.ccell.2015.04.006 From NLM Medline. Rahman, M. A.; Lin, K. T.; Bradley, R. K.; Abdel-Wahab, O.; Krainer, A. R. Recurrent SRSF2 mutations in MDS affect both splicing and NMD. Genes Dev 2020, 34 (5-6), 413–427. DOI: 10.1101/gad.332270.119 From NLM Medline. Zhang, J.; Lieu, Y. K.; Ali, A. M.; Penson, A.; Reggio, K. S.; Rabadan, R.; Raza, A.; Mukherjee, S.; Manley, J. L. Disease-associated mutation in SRSF2 misregulates splicing by altering RNA-binding affinities. Proc Natl Acad Sci U S A 2015, 112 (34), E4726–4734. DOI: 10.1073/pnas.1514105112 From NLM Medline.

(39) van der Feltz, C.; Hoskins, A. A. Structural and functional modularity of the U2 snRNP in pre-mRNA splicing. Crit Rev Biochem Mol Biol 2019, 54 (5), 443–465. DOI: 10.1080/10409238.2019.1691497 From NLM Medline.

(40) Staley, J. P.; Guthrie, C. Mechanical devices of the spliceosome: motors, clocks, springs, and things. Cell 1998, 92 (3), 315–326. DOI: 10.1016/s0092-8674(00)80925-3 From NLM Medline.

