## Supplemental Information for "A Yeast-Based High-Throughput Screening Platform for the Discovery of Novel pre-mRNA Splicing Modulators"

**Supplemental Materials:**

Supplemental Figures S1-S3
Supplemental Tables S1-S3

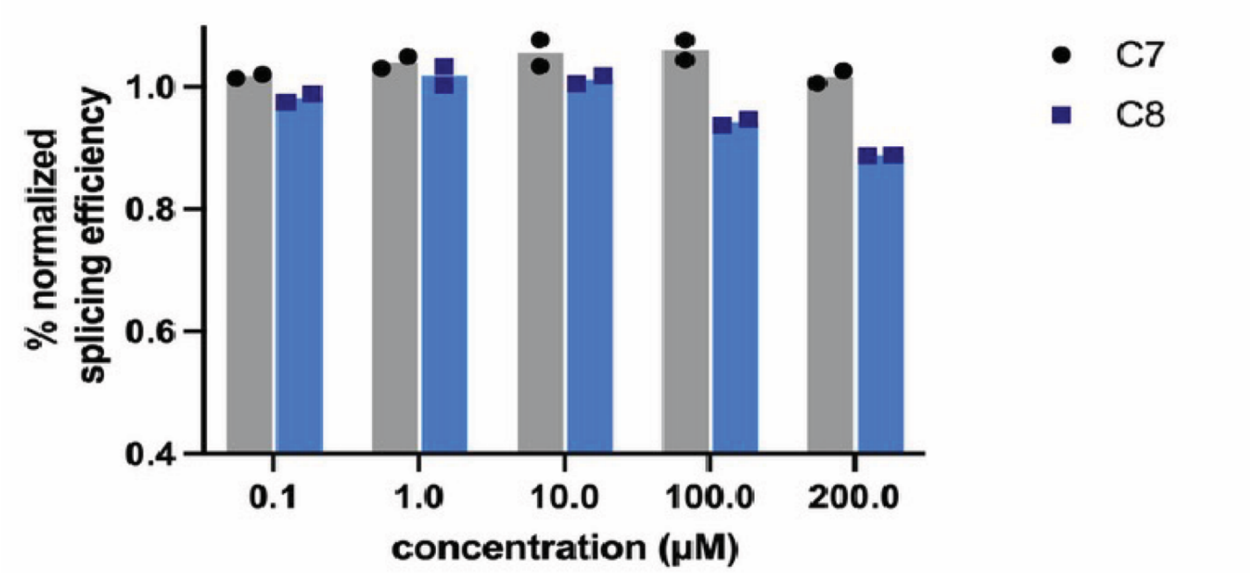


**Figure S1. Compound C7 Does Not Reduce Splicing Efficiency in HeLa Nuclear Extract**
*In vitro* splicing assay results using HeLa nuclear extracts and the AdML pre-mRNA substrate in the presence of compounds C7 and C8. Bar heights represent the averages of two technical replicates (dots, squares).

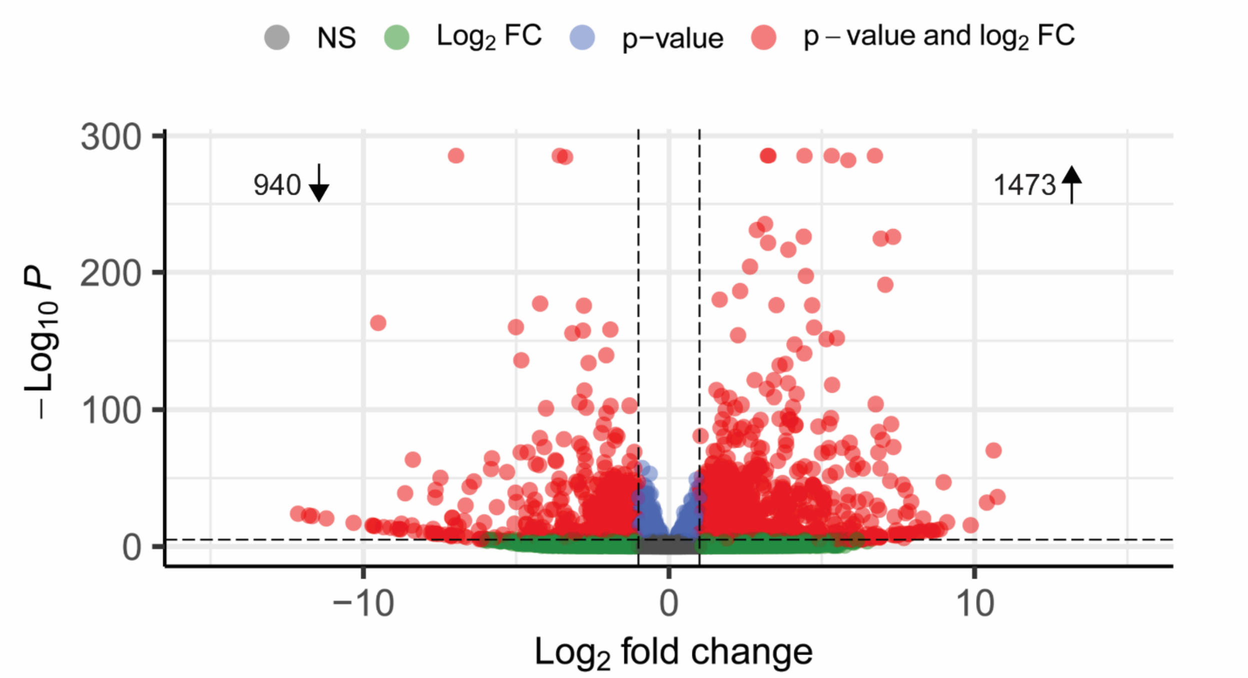


**Figure S2. DEGs in P95H Mutant Cells relative to WT upon DMSO treatment**
Differential gene analysis of $SRSF2^{WT/WT}$ and $SRSF2^{WT/P95H}$ mutant cells in the DMSO control. DEGs were defined as those with $\log_2$ fold change <-1 or >1 and a $-\log_{10}$ *p*-value >0.05 (red dots). Genes downregulated in the P95H cell line are shown on the left, and upregulated genes on the right.

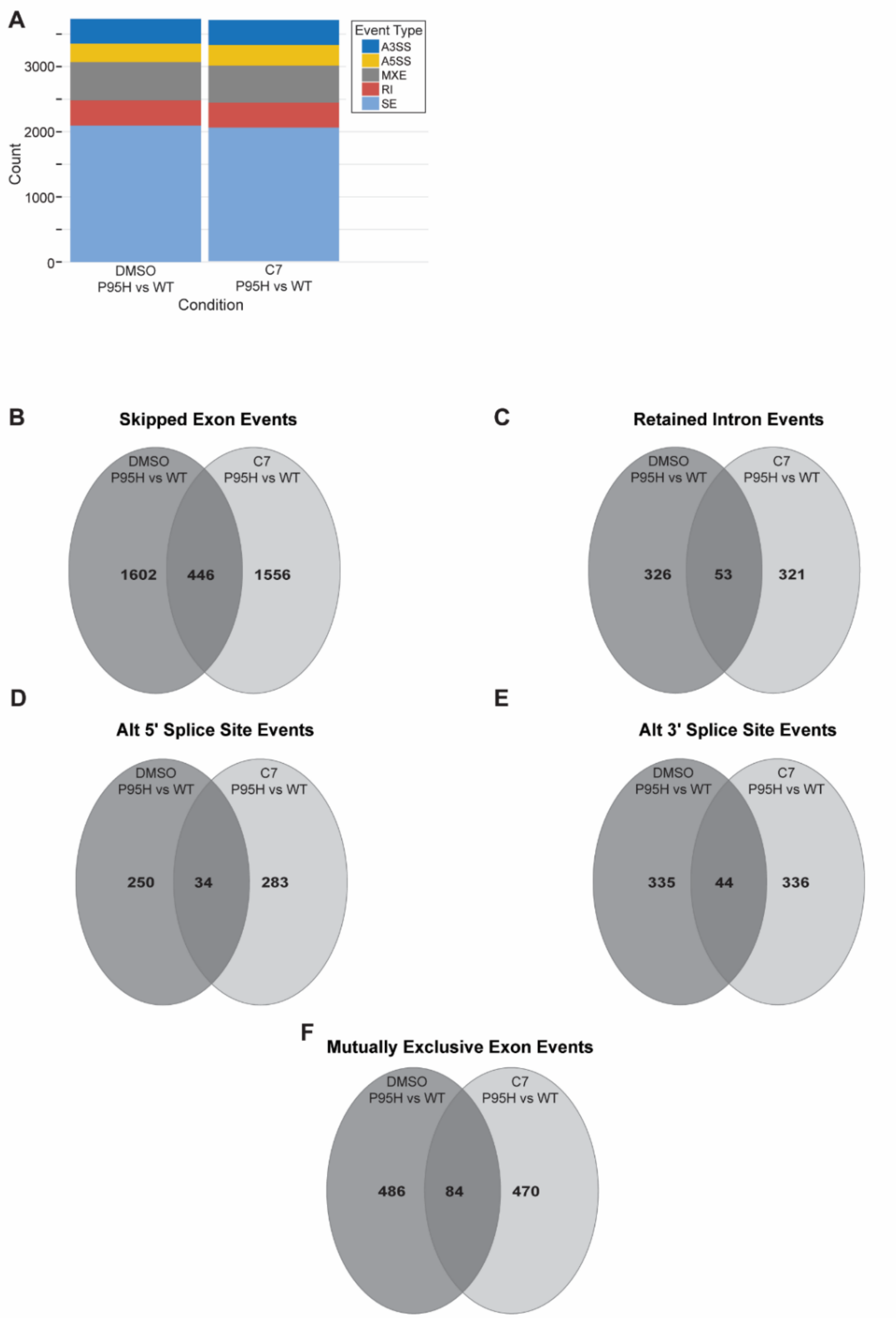


**Figure S3. C7 Treatment Induces Changes in pre-mRNA Splicing (A)** Stacked bar graph showing the distribution of novel alternative splicing events found in P95H mutant cells relative to WT after treatment with DMSO (left bar) and compound C7 (right bar). A3SS/A5SS: alternative 3'/5' splice sites; MXE: mutually exclusive exons; RI: retained introns; SE: skipped exons. **(B–F)** Venn diagrams showing unique splicing events in the P95H mutant cells in comparison to WT upon treatment with DMSO (dark gray) or compound C7 (light gray). Overlapping events were unique to the P95H cell line relative to WT and found upon treatment with either DMSO or C7.

**Table S1. Selleck FDA Pilot Screening Hits**

| Compound | Use/MoA | Specificity |
|---|---|---|
| Benazepril | Hypertension treatment/ACE inhibitor | Hsh155 |
| Bifonazole | Antifungal | Hsh155 |
| Butoconazole | Antifungal | Both |
| Caspofungin | Antifungal | Both |
| Climbazole | Antifungal | Both |
| Cyproheptadine | Antihistamine | Both |
| Cytidine | Antidepressant | Hs-Hsh155 |
| Econazole | Antifungal | Both |
| Everolimus | Immunosuppressant | Both |
| Fenticonazole | Antifungal | Both |
| Isocinazole | Antifungal | Both |
| Miconazole | Antifungal | Both |
| Rapaycin | Antibiotic/immunosupressant | Both |
| Rivastigmine | cognition enhancing med | Both |
| Sulconazole | Antifungal | Both |
| Tioconazole | Antifungal | Hsh155 |
| Voriconazole | Antifungal | Both |
| Amorolfine | Antifungal | Hs-Hsh155 |

### Table S2. Secondary Screening Hits

| Structure | Name | Library | Specificiy | Lipinski Rules | PAINS parameters |
|---|---|---|---|---|---|
| 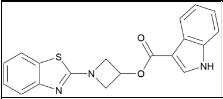 | 1-(1,3-benzothiazol-2-yl)azetidin-3-yl 1H-indole-3-carboxylate | Life Chemicals 4 | WTBrr2, HsBrr2 | Pass | Pass |
| 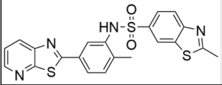 | 2-methyl-N-(2-methyl-5-{[1,3]thiazolo[5,4-b]pyridin-2-yl}phenyl)-1,3-benzothiazole-6-sulfonamide | Life Chemicals 4 | WTHsh155 | Pass | Pass |
| 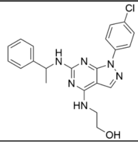 | 2-[[1-(4-chlorophenyl)-6-[[(1S)-1-phenylethyl]amino]pyrazolo[3,4-d]pyrimidin-4-yl]amino]ethanol | Life Chemicals 4 | HsBrr2 | Pass | Pass |
| 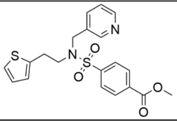 | methyl 4-{[(pyridin-3-yl)methyl][2-(thiophen-2-yl)ethyl]sulfamoyl}benzoate | Life Chemicals 4 | WTBrr2 | Pass | Pass |
| 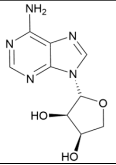 | 9-beta-d-erythrofuranosyladenine (EFA) | NCI:DTP | Nonspecific | Pass | Pass |
| 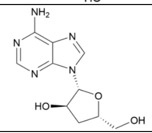 | 3'-Deoxyadenosine (Cordycepin) | NCI:DTP | Nonspecific | Pass | Pass |
| 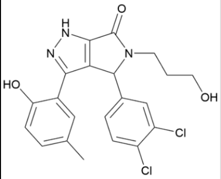 | 4-(3,4-dichlorophenyl)-3-(2-hydroxy-3,5-dimethylphenyl)-5-(3-hydroxypropyl)-1,4-dihydropyrrolo[3,4-c]pyrazol-6-one | NCI:NExT | HsBrr2 | Pass | Pass |
| 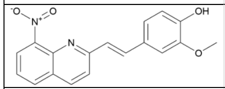 | 2-methoxy-4-[(E)-2-(8-nitroquinolin-2-yl)ethenyl]phenol (drug C8) | NCI:NExT | HsBrr2 | Pass | Pass |
| 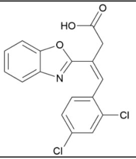 | (Z)-3-(1,3-benzoxazol-2-yl)-4-(2,4-dichlorophenyl)but-3-enoic acid | NCI:NExT | HsBrr2 | Pass | Pass |
| 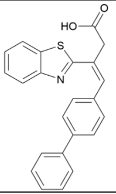 | (Z)-3-(1,3-benzothiazol-2-yl)-4-(4-phenylphenyl)but-3-enoic acid | NCI:NExT | WTHsh155 | Pass | Pass |
| 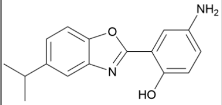 | 4-Amino-2-[5-(propan-2-yl)-1,3-benzoxazol-2-yl]phenol (drug C7) | NCI:NExT | HsBrr2 | Pass | Pass |

### Table S3. GO-term Enrichment of Alternative Splicing Events Unique to P95H Mutant Cells Treated with C7 relative to WT

| Alt Splicing Type | Enrichment FDR | nGenes | Fold Enrichment | Pathway |
|---|---|---|---|---|
| A3SS | 0.00515325 | 10 | 6.45954492 | GO:0002181 cytoplasmic translation |
| A3SS | 0.01622916 | 20 | 2.9331607 | GO:0006412 translation |
| A3SS | 0.01729273 | 20 | 2.82505896 | GO:0043043 peptide biosynthetic proc. |
| A3SS | 0.01729273 | 22 | 2.63770364 | GO:0043604 amide biosynthetic proc. |
| A3SS | 0.02419751 | 22 | 2.53208988 | GO:0006518 peptide metabolic proc. |
| A3SS | 0.02441377 | 23 | 2.44411986 | GO:0006396 RNA processing |
| A3SS | 0.01729273 | 26 | 2.37448561 | GO:0034645 cellular macromolecule biosynthetic proc. |
| A3SS | 0.00515325 | 37 | 2.25331371 | GO:1901566 organonitrogen compound biosynthetic proc. |
| A5SS | 0.024031898 | 3 | 29.35970915 | GO:0006287 base-excision repair gap-filling |
| A5SS | 0.024031898 | 3 | 29.35970915 | GO:1902969 mitotic DNA replication |
| A5SS | 0.00897284 | 4 | 23.82816975 | GO:0006268 DNA unwinding involved in DNA replication |
| A5SS | 0.028084961 | 4 | 14.42231327 | GO:0033260 nuclear DNA replication |
| A5SS | 0.032995488 | 4 | 13.04875962 | GO:0044786 cell cycle DNA replication |
| A5SS | 0.023896835 | 5 | 10.87396635 | GO:0042255 ribosome assembly |
| A5SS | 0.029093228 | 6 | 7.27497218 | GO:0071103 DNA conformation change |
| A5SS | 0.00897284 | 8 | 6.683511027 | GO:0006261 DNA-templated DNA replication |
| A5SS | 0.00897284 | 11 | 4.608965555 | GO:0042254 ribosome biogenesis |
| A5SS | 0.032995488 | 10 | 3.937125749 | GO:0006310 DNA recombination |
| A5SS | 0.001823076 | 18 | 3.800023989 | GO:0006281 DNA repair |
| A5SS | 0.00897284 | 14 | 3.768502288 | GO:0022613 ribonucleoprotein complex biogenesis |
| A5SS | 0.032995488 | 12 | 3.40402425 | GO:0034470 ncRNA processing |
| A5SS | 0.021842731 | 15 | 3.186325024 | GO:0034660 ncRNA metabolic proc. |
| A5SS | 0.000614589 | 26 | 3.130326342 | GO:0006259 DNA metabolic proc. |
| A5SS | 0.003695002 | 23 | 2.912454204 | GO:0006396 RNA processing |
| A5SS | 0.008035651 | 21 | 2.900455138 | GO:0006974 cellular response to DNA damage stimulus |
| A5SS | 0.024031898 | 19 | 2.613682274 | GO:0000278 mitotic cell cycle |
| A5SS | 0.024031898 | 21 | 2.452899827 | GO:0051276 chromosome organization |
| A5SS | 0.00897284 | 53 | 1.72649423 | GO:0006996 organelle organization |
| RI | 0.00976974 | 64 | 1.67575291 | Cellular protein modification process |
| RI | 0.00976974 | 31 | 2.33850003 | Cell cycle process |
| RI | 0.00976974 | 64 | 1.67575291 | Protein modification process |
| RI | 0.01172098 | 26 | 2.51641391 | Mitotic cell cycle |
| RI | 0.01445863 | 36 | 2.0343783 | Cell cycle |
| RI | 0.01445863 | 65 | 1.61216407 | Macromolecule modification |
| RI | 0.01445863 | 55 | 1.69585565 | Cellular component biogenesis |
| RI | 0.01445863 | 8 | 6.7374021 | Positive regulation of proteasomal protein catabolic process |
| RI | 0.01445863 | 23 | 2.542198 | Mitotic cell cycle process |
| RI | 0.01752097 | 51 | 1.71957746 | Cellular component assembly |
| RI | 0.01967816 | 2 | 112.851485 | Neural plate elongation |
| RI | 0.01967816 | 2 | 112.851485 | Convergent extension involved in neural plate elongation |
| RI | 0.01967816 | 8 | 5.82459278 | Positive regulation of proteolysis involved in cellular protein catabolic process |
| RI | 0.02501625 | 13 | 3.47646755 | Positive regulation of proteolysis |
| RI | 0.04097803 | 8 | 5.1006321 | Positive regulation of cellular protein catabolic process |
| RI | 0.04143575 | 34 | 1.90042125 | Phosphorylation |
| RI | 0.04440153 | 3 | 24.1824611 | CTP biosynthetic process |
| RI | 0.04440153 | 16 | 2.7693616 | Regulation of mitotic cell cycle |
| RI | 0.04445504 | 3 | 22.570297 | Fructose metabolic process |
| RI | 0.04445504 | 46 | 1.66543738 | Phosphate-containing compound metabolic process |
| SE | 0.032711726 | 3 | 23.1902027 | GO:1901483 reg. of transcription factor catabolic proc. |
| SE | 0.026891774 | 25 | 2.321341612 | GO:0048193 Golgi vesicle transport |
| SE | 0.023895547 | 26 | 2.310135135 | GO:0006310 DNA recombination |
| SE | 0.017804515 | 33 | 2.11256505 | GO:0034470 ncRNA processing |
| SE | 0.010673076 | 42 | 2.013412948 | GO:0034660 ncRNA metabolic proc. |
| SE | 0.032711726 | 47 | 1.765799153 | GO:0016071 mRNA metabolic proc. |
| SE | 0.010811217 | 61 | 1.743194535 | GO:0006396 RNA processing |
| SE | 0.017804515 | 62 | 1.684584145 | GO:0006259 DNA metabolic proc. |
| SE | 0.010673076 | 87 | 1.574057059 | GO:0046907 intracellular transport |
| SE | 0.024212078 | 91 | 1.490330823 | GO:1901566 organonitrogen compound biosynthetic proc. |
| SE | 1.38E-05 | 200 | 1.470293403 | GO:0006996 organelle organization |
| SE | 0.017804515 | 103 | 1.46427027 | GO:0071705 nitrogen compound transport |
| SE | 0.017804515 | 120 | 1.417277476 | GO:0071702 organic substance transport |
| SE | 0.00089922 | 201 | 1.380185282 | GO:0018130 heterocycle biosynthetic proc. |
| SE | 0.001275314 | 199 | 1.363424282 | GO:0019438 aromatic compound biosynthetic proc. |
| SE | 0.001058461 | 206 | 1.362184704 | GO:1901362 organic cyclic compound biosynthetic proc. |
| SE | 0.002211429 | 194 | 1.35560055 | GO:0034654 nucleobase-containing compound biosynthetic proc. |
| SE | 0.010673076 | 191 | 1.317664351 | GO:0031326 reg. of cellular biosynthetic proc. |
| SE | 0.016957976 | 183 | 1.313160701 | GO:0010556 reg. of macromolecule biosynthetic proc. |
